# Potential impact on coagulopathy of gene variants of coagulation related proteins that interact with SARS-CoV-2

**DOI:** 10.1101/2020.09.08.272328

**Authors:** David Holcomb, Aikaterini Alexaki, Nancy Hernandez, Kyle Laurie, Jacob Kames, Nobuko Hamasaki-Katagiri, Anton A. Komar, Michael DiCuccio, Chava Kimchi-Sarfaty

**Author notes:** Equal contribution (David Holcomb, Aikaterini Alexaki, Nancy Hernandez). Corresponding authors: Chava Kimchi-Sarfaty, PhD; Division of Plasma Protein Therapeutics, Office of Tissue and Advanced Therapies, Center for Biologics Evaluation and Research, Food and Drug Administration, Silver Spring, USA, Michael DiCuccio, MD; National Center of Biotechnology Information, National Institutes of Health, Bethesda, MD, USA. Author contribution DH created software, as well as investigation and formal analysis. AA wrote original draft, as well as further review and editing. NH performed investigation and visualization. KL assisted in investigation. JK, NHK, AAK, MD, and CKS provided conceptualization, and reviewed and edited the manuscript. CKS was also involved in producing methodology and administration of the project.

## Abstract

Thrombosis has been one of the complications of the Coronavirus disease of 2019 (COVID-19), often associated with poor prognosis. There is a well-recognized link between coagulation and inflammation, however, the extent of thrombotic events associated with COVID-19 warrants further investigation. Poly(A) Binding Protein Cytoplasmic 4 (PABPC4), Serine/Cysteine Proteinase Inhibitor Clade G Member 1 (SERPING1) and Vitamin K epOxide Reductase Complex subunit 1 (VKORC1), which are all proteins linked to coagulation, have been shown to interact with SARS proteins. We computationally examined the interaction of these with SARS-CoV-2 proteins and, in the case of VKORC1, we describe its binding to ORF7a in detail. We examined the occurrence of variants of each of these proteins across populations and interrogated their potential contribution to COVID-19 severity. Potential mechanisms by which some of these variants may contribute to disease are proposed. Some of these variants are prevalent in minority groups that are disproportionally affected by severe COVID-19. Therefore, we are proposing that further investigation around these variants may lead to better understanding of disease pathogenesis in minority groups and more informed therapeutic approaches.

**Author summary:** Increased blood clotting, especially in the lungs, is a common complication of COVID-19. Infectious diseases cause inflammation which in turn can contribute to increased blood clotting. However, the extent of clot formation that is seen in the lungs of COVID-19 patients suggests that there may be a more direct link. We identified three human proteins that are involved indirectly in the blood clotting cascade and have been shown to interact with proteins of SARS virus, which is closely related to the novel coronavirus. We examined computationally the interaction of these human proteins with the viral proteins. We looked for genetic variants of these proteins and examined how these variants are distributed across populations. We investigated whether variants of these genes could impact severity of COVID-19. Further investigation around these variants may provide clues for the pathogenesis of COVID-19 particularly in minority groups.

## Introduction

The Coronavirus disease of 2019 (COVID-19) has been associated with coagulopathy, particularly microclots in the lungs [1] [2] [3] [4] [5], that correlates with disease severity [6] [7] [8] [9]. There is extensive cross-talk between inflammation and coagulation, and inflammation is presumed to have a role in the observed coagulation phenotype. However, the widespread thrombotic events that are seen in severe COVID-19 patients suggest that there may be a more direct link.

In a study conducted before the onset of the COVID-19 pandemic, the severe acute respiratory syndrome (SARS) coronavirus (CoV)-host interactome was investigated. A few proteins related to the coagulation cascade were experimentally identified to interact with viral proteins. Poly(A) Binding Protein Cytoplasmic 4 (PABPC4) was shown to interact with the nucleocapsid (N) protein. Serine/Cysteine Proteinase Inhibitor Clade G Member 1 (SERPING1 or C1 inhibitor) was shown to interact with nsp14, ORF14, ORF3b, ORF7b, nsp2, nsp8 and nsp13. In addition, Vitamin K epOxide Reductase Complex subunit 1 (VKORC1) was shown to interact with the SARS protein ORF7a. The interactions were initially identified by a high-throughput yeast two-hybrid system and confirmed with LUMIER assay [10].

PABPC4 localizes primarily to the cytoplasm and binds to the poly(A) tail present at the 3-prime end of mRNA. However, it is also found in the surface of thrombin-activated platelets, and therefore it is known as activated-platelet protein-1 (APP-1) [11] [12]. PABPC4 may also be involved in the regulation of protein translation in platelets and megakaryocytes or may participate in the binding or stabilization of polyadenylates in platelet dense granules [13]. SERPING1 is a plasma protease involved in the complement, intrinsic coagulation and fibrinolytic pathways. In the coagulation cascade, SERPING1 inactivates plasma kallikrein, factor XIIa and factor XIIf. The absence of sufficient levels of functional SERPING1 leads to hereditary angioedema (HAE), which is mediated by sustained activation of kallikrein leading to cleavage of high molecular weight kininogen (HMWK), producing bradykinin [14].

VKORC1 is an enzyme critical for coagulation due to its role in converting vitamin K epoxide into active vitamin K [15], the rate-limiting step in the physiological process of vitamin K recycling. Importantly, vitamin K is necessary for the carboxylation of glutamic acid residues to produce Gla residues. Several human proteins have domains with Gla residues, including coagulation factors II, VII, IX, X, and anticoagulant proteins C, S, and Z. VKORC1 is expressed in all tissues, but particularly in the liver, lungs, and female reproductive system. It is generally embedded in the endoplasmic reticulum [16].

Dietary vitamin K deficiency is associated with coagulopathy, specifically bleeding. Vitamin K antagonists are anticoagulant drugs that work by inhibiting the activity of VKORC1, reducing the levels of available active vitamin K and coagulation factors. Of the vitamin K antagonists, warfarin is most commonly used. Some variants in *VKORC1*, particularly those common in African and African American populations, are reported to result in warfarin resistance. Warfarin response is also dependent on dietary factors and liver function [17]. For these reasons, dosing warfarin is complicated, and genotyping of *VKORC1* to determine the presence of known polymorphisms (such as c.1173C>T) is recommended before initiating warfarin treatment. The impact of viral protein interactions with VKORC1, SERPING1 and PABPC4 on patient outcomes in COVID-19 infection is unknown. While comorbidities, age, and other factors will impact the predisposition to thrombosis or coagulopathy, binding of viral proteins to coagulation related proteins may be partially responsible for the prothrombotic phenotype that is seen in COVID-19 patients.

Through computational modeling, we examined the binding of VKORC1, SERPING1 and PABPC4 to SARS-CoV-2 proteins and generated additional evidence for the binding of ORF7a to VKORC1. We analyzed COVID-19 genome-wide association study (GWAS) results to find the most influential variants from these genes and characterize them to find potential means of effect. Moreover, we investigated several *VKORC1, SERPING1* and *PABPC4* variants that may be associated with coagulopathy and we identified some *VKORC1* variants that may result in warfarin resistance. In particular, we highlight two variants, which are enriched in certain ethnic groups. Better understanding of the contribution of these genes and their variants to COVID-19 pathogenesis may lead to new therapeutic avenues and improved prognosis. This may be of crucial importance for minority groups that are disproportionally affected by severe COVID-19.

## Methods

### Structural similarities and computational docking of proteins

To assess the binding of SARS-CoV-2 ORF7a and human VKORC1, we used I-TASSER [18] [19] [20] to generate homology models for both proteins. Using the models with the best C-scores from I-TASSER, we then used Zdock [21] to find potential binding sites. From Zdock, we took the five top scoring protein-protein complexes as input to Rosetta Prepack and Rosetta Dock [22] [23] [24] [25] to further refine the models by using rigid body perturbations. Finally, the top model was further refined through Rosetta Dock local refinement.

In addition, to verify the binding of PABPC4 and SERPING1 with SARS-CoV-2 proteins, we created homology models for each using I-TASSER and Robetta [26] [27]. However, due to the lack of structural data from similar proteins, sections of the models for PABPC4 and SERPING1 were of low quality. For this reason, we used Blast and Clustal Omega to create multiple sequence alignments (MSAs) of proteins similar to interacting SARS proteins, and computed the percent of columns of the homologous SARS-CoV-2 protein matching the SARS protein, as well as a loglikelihood score to measure the probability that the SARS-CoV-2 homolog would be included in the MSA (Table 1). In addition, the MSAs were filtered to remove duplicate sequences by performing affinity propagation clustering with the Levenshtein distance matrix formed from the sequences. Only the cluster centers, SARS, and SARS-CoV-2 sequences were used in the MSA. This was done to account for the large number of very similar sequences, generally from different strains of SARS-CoV-2.

**Table 1:** Sequence homology of selected SARS and SARS-CoV-2 proteins.

| Protein | Fraction Matching | Loglikelihood |
| --- | --- | --- |
| N | 0.888626 | -0.04824 |
| ORF7a | 0.827869 | -0.05688 |
| nsp14 | 0.884393 | -0.19572 |
| ORF7b | 0.795455 | -0.01103 |
| nsp3 | 0.75078 | -0.46736 |
| nsp2 | 0.681818 | -0.44764 |
| nsp8 | 0.969697 | -0.12897 |
| nsp13 | 0.948767 | -0.1215 |

MSA fraction matching is the fraction of positions in the SARS-CoV-2 protein matching the homologous SARS protein, when both are aligned in an MSA. Higher number indicates more conserved position and the range is between 0 and 1.
MSA likelihood is the fraction of sequences in an MSA matching SARS-CoV-2 for a given column. Assuming all columns are independent, $\prod_i P(x_i)$ gives the probability of finding the SARS-CoV-2 sequence in the MSA sequences, which ranges between 0 and 1. Taking log of this value gives $\log(\prod_i P(x_i)) = \sum_i (\log P(x_i))$ , an additive loglikelihood score which is nonpositive, with lower values indicating more positions in the SARS-CoV-2 sequence that differ from the MSA sequences.

We used the ORF7a homology model to query Dali [28] for similar protein structures. The top structures in sequence and structural similarity were the ORF7a proteins for SARS and SARS-CoV-2 (PDBs 1YO4 and 6W37). All human proteins interacting with VKORC1 were taken from BIOGRID, the Biological General Repository for Interaction Datasets [29] [30]. In addition, we queried Dali against all other viral protein structures, as modeled in I-TASSER.

### Relevant variants from COVID19 HGI GWAS metastudies

All variants from the genomic region containing *VKORC1, SERPING1*, and *PABPC4* ±6000 bp were taken from the ANA2, ANA5, and ANA7 metastudies from COVID19 Host Genetics Initiative [31] and The Severe Covid-19 GWAS Group [32] (Supplementary Table S1 and Table 2). We filtered the resulting variants to keep only those with metastudy p-value below 0.05. The resulting variants were all in non-coding regions, therefore, amino acid and codon features do not apply.

**Table 2:** Possible predicted effect of variants in VKORC, SERPING1 and PABPC4.

| Transcript | Location | Fraction matching in MSA | Change in splicing | Average change in mRNA MFE (Z-score) | miRNA summary |
| --- | --- | --- | --- | --- | --- |
| VKORC1 |  |  |  |  |  |
| NM_024006.4:c.-4931C>T | 5' UTR | 0.431818 |  | 2.20025 |  |
| NM_024006.4:c.-4851C>T | 5' UTR | 0.754967 |  | 1.06847 |  |
| NM_024006.4:c.-2834C>A | 5' UTR | <u>0.950943</u> |  |  | <u>miRNA gained</u> |
| NM_024006.4:c.-1639G>A | 5' UTR | 0.0625 | Possible splicing change |  |  |
| NM_024006.4:c.174-136C>T | Intron | 0.020228 |  | 0.83549 |  |
| NM_024006.4:c.283+124G>C | Intron | 0.133333 |  | -1.44871 |  |
| NM_024006.4:c.283+837T>C | Intron | 0.146727 |  | 1.35367 |  |
| SERPING1 |  |  |  |  |  |
| NM_000062.2:c.-3537C>G | 5' UTR | 0.009615 |  |  | <u>miRNA decrease</u> |
| NM_000062.2:c.-2415G>A | 5' UTR | 0.033708 | Possible splicing change | 0.77565 |  |
| NM_000062.2:c.-1675G>A | 5' UTR | 0.426901 |  |  | <u>miRNA gained</u> |
| NM_000062.2:c.52-696C>T | Intron | 0.068027 | <u>Likely splicing change</u> | 0.30003 |  |
| NM_000062.2:c.52-130C>T | Intron | 0.833333 |  |  |  |
| NM_000062.2:c.52-130C>T | Intron | 0.833333 |  | -0.28155 |  |
| NM_000062.2:c.550+794C>A | Intron | 0.693694 |  | 0.71906 |  |
| NM_000062.2:c.685+88G>A | Intron | 0.769231 |  | -0.75902 |  |
| <u>NM_000062.2:c.685+659C>T</u> | Intron | 0.581818 |  | -0.47852 | miRNA decrease |
| <u>NM_000062.2:c.685+659C>T</u> | Intron | 0.679245 |  | -0.68662 |  |
| <u>NM_000062.2:c.685+1100C>T</u> | Intron | 0.772313 |  | <u>2.01226</u> |  |
| <u>NM_000062.2:c.685+1391C>T</u> | Intron | 0.841912 | <u>Likely splicing change</u> |  |  |
| <u>NM_000062.2:c.685+1550G>T</u> | Intron | <u>0.926641</u> |  |  |  |
| <u>NM_000062.2:c.685+1770C>T</u> | Intron | 0.793594 |  | 0.84106 |  |
| <u>NM_000062.2:c.1029+926G>T</u> | Intron | 0.015723 |  |  | miRNA gained |
| NM_000062.2:c.1029+1443G>C | Intron | 0.595745 |  |  | miRNA gained |
| <u>NM_000062.2:c.1029+2110T>C</u> | Intron | 0.393617 | <u>Likely splicing change</u> |  |  |
| NM_000062.2:c.1029+2111G>A | Intron | 0.687117 |  |  | miRNA gained |
| NM_000062.2:c.1030-2243T>G | Intron | 0.026616 |  | -1.68665 |  |
| NM_000062.2:c.1030-1975G>C | Intron | 0.02551 |  | <u>1.92651</u> |  |
| NM_000062.2:c.1030-1436T>C | Intron | 0.823529 |  |  |  |
| NM_000062.2:c.1030-20A>G | Intron | 0.25 |  | -0.76751 |  |
| NM_000062.2:c.1438G>A | Exon | 0.399177 | Possible<br>splicing<br>change |  |  |
| NM_000062.2:c.*1323G>A | 3' UTR | 0.878788 |  |  | miRNA lost |
| NM_000062.2:c.*1521G>T | 3' UTR | 0.65873 |  |  |  |
| NM_000062.2:c.*2614A>T | 3' UTR | 0.016129 |  |  | miRNA gained |
| PABPC4 |  |  |  |  |  |
| NM_003819.3:c.-5600T>C | 5' UTR | 0.48 |  |  | <u>miRNA lost</u> |
| NM_003819.3:c.-4432G>A | 5' UTR | 0.009317 |  |  | <u>miRNA gained</u> |
| NM_003819.3:c.-4428A>G | 5' UTR | 0.221505 |  |  |  |
| NM_003819.3:c.-3677T>G | 5' UTR | 0.021645 | Possible<br>splicing<br>change | -0.59763 |  |
| NM_003819.3:c.-3636G>A | 5' UTR | 0.022378 |  | 2.31727 |  |
| NM_003819.3:c.-3198T>C | 5' UTR | 0.856079 | Possible<br>splicing<br>change |  | <u>miRNA lost</u> |
| NM_003819.3:c.-2286T>G | 5' UTR | 0.210526 |  |  |  |
| NM_003819.3:c.-650C>T | 5' UTR | 0.829978 |  |  | <u>miRNA lost</u> |
| NM_003819.3:c.193+796C>G | Intron | 0.666667 |  |  |  |
| NM_003819.3:c.504-254C>A | Intron | 0.247191 |  | 0.26322 |  |
| NM_003819.3:c.738+85T>C | Intron | 0.333333 |  |  | miRNA lost |
| NM_003819.3:c.877-387C>T | Intron | <u>1</u> |  |  | miRNA lost |
| NM_003819.3:c.972+53A>T | Intron | <u>1</u> |  |  |  |
| NM_003819.3:c.972+704C>G | Intron | 0.5 |  |  |  |
| NM_003819.3:c.1333+26C>G | Intron | 0.3125 | <u>Likely</u><br><u>splicing</u><br><u>change</u> |  |  |
| NM_003819.3:c.1621-348C>G | Intron | <u>1</u> |  |  |  |
| NM_003819.3:c.*765C>A | 3' UTR | <u>1</u> |  | <u>1.96974</u> |  |
| NM_003819.3:c.*1261C>T | 3' UTR | 0.771242 |  | -0.91257 | miRNA decrease |
| NM_003819.3:c.*4685A>G | 3' UTR | 0.054945 |  | -3.4524 | miRNA decrease |
| NM_003819.3:c.*5316C>T | 3' UTR | 0.696181 | Possible<br>splicing<br>change |  | miRNA lost |

Change in splicing is presented when all tools find a change in splicing and all hexamer scores are greater than one standard deviation from the mean, and is marked in red when the variant appears in an intron. mRNA MFE changes are normalized (converted into a Z-score) for KineFold, remuRNA, and mFold, then averaged. When all three mRNA MFE changes are above one standard deviation, we mark the value in underline. miRNA summaries are presented when all miRNA changes agree in direction, and the total change is at least 5. miRNA changes are underlined when the variant appears upstream.

We characterized these variants in terms of splicing, using hexamer scoring tools [33] [34], ESEfinder [35] [36], ExonScan [37] [38] [39], and FAS-ESS [37]. Where ESEfinder, ExonScan, and FAS-ESS found a change in splicing potential between the wild type (WT) and mutant, the change was reported in Table 2 as “Change in splicing”. When the variant occurred in an intron as opposed to a UTR, we further highlighted the value.

Then, we calculated mRNA mean free energy using Kinefold [40], mFold [41] [42] [43], and remuRNA [44]. When all three tools were in agreement regarding the direction of the change, the changes in mRNA MFE were converted into Z-scores using mean and standard deviation values computed by randomly sampling WT and mutant sequences. The average of the three Z-scores is reported in Table 2 as “Average change in mRNA MFE (Z-score)”.

We also analyzed miRNA binding changes using miRDB [45] [46]. For any variant, there may be multiple affected miRNA species. miRNA binding scores are provided for both the WT and mutant flanking 501 nucleotides in Supplementary Table S2, and a summary of miRNA binding changes is provided in Table 2. When all miRNA binding changes were in the same direction, we summarized the effect.

We analyzed conservation using fraction matching in a nucleotide MSA, computed as the fraction of sequences in the MSA matching the wild type sequence in the appropriate column. This value is included in Table 2 as “Fraction matching in MSA”.

Finally, we collected population prevalence data from dbSNP (Supplementary Table S2 and Table 3).

**Table 3:** Population frequencies (gnomAD) of GWAS variants of VKORC1, SERPING1, and PABPC4.

| Transcript | Global | African | American | Ashkenazi Jewish | East Asian | European | Other |
| --- | --- | --- | --- | --- | --- | --- | --- |
| VKORC1 |  |  |  |  |  |  |  |
| NM_024006.4:c.-4931C>T | 0.5758 | 0.5408 | 0.542 | 0.514 | <u>0.1007</u> | 0.63146 | 0.619 |
| NM_024006.4:c.-4851C>T |  |  |  |  |  |  |  |
| NM_024006.4:c.-2834C>A | 0.0049 | 0.0001 | 0.001 | 0 | 0 | 0.00762 | 0.007 |
| NM_024006.4:c.-1639G>A | 0.326 | <u>0.1009</u> | 0.444 | 0.476 | <u>0.8996</u> | 0.37236 | 0.369 |
| NM_024006.4:c.174-136C>T | 0.3261 | <u>0.1009</u> | 0.443 | 0.476 | <u>0.8995</u> | 0.37264 | 0.37 |
| NM_024006.4:c.283+124G>C | 0.4163 | 0.2564 | 0.442 |  | <u>0.8849</u> |  |  |
| NM_024006.4:c.283+837T>C | 0.6431 | 0.7907 | 0.546 | 0.517 | <u>0.1017</u> | 0.62682 | 0.628 |
| NM_014699.3:c.*2082G>C | 0.0049 | 0.0001 | 0.001 | 0 | 0 | 0.00763 | 0.007 |
| NM_014699.3:c.*2737G>T | 0.0048 | 0.017 | 0.002 | 0 | 0 | 0.00005 | 0 |
| SERPING1 |  |  |  |  |  |  |  |
| NM_000062.2:c.-3537C>G | 0.0275 | 0.0078 | 0.021 | 0.017 | 0 | 0.03846 | 0.041 |
| NM_000062.2:c.-2415G>A | 0.0939 | 0.0866 | 0.059 | 0.141 | 0.1113 | 0.09708 | 0.087 |
| NM_000062.2:c.-1675G>A | 0.0937 | 0.0862 | 0.059 | 0.141 | 0.1105 | 0.09697 | 0.087 |
| NM_000062.2:c.52-696C>T | 0.3927 | 0.4767 | 0.529 | 0.452 | 0.7655 | 0.31673 | 0.386 |
| NM_000062.2:c.52-130C>T | 0.385 | 0.448 | 0.525 | 0.455 | 0.7668 | 0.31739 | 0.385 |
| NM_000062.2:c.52-130C>T | 0.385 | 0.448 | 0.525 | 0.455 | 0.7668 | 0.31739 | 0.385 |
| NM_000062.2:c.550+794C>A | 0.3936 | 0.4761 | 0.531 | 0.451 | 0.7697 | 0.31779 | 0.388 |
| NM_000062.2:c.685+88G>A | 0.2225 | 0.1006 | 0.15 | 0.262 | 0.1157 | 0.28733 | 0.273 |
| NM_000062.2:c.685+1391C>T | 0.0248 | 0.0059 | 0.022 | 0.035 | 0 | 0.03499 | 0.038 |
| NM_000062.2:c.685+659C>T | 0.3901 | 0.4743 | 0.541 | 0.455 | 0.769 | 0.31084 | 0.381 |
| NM_000062.2:c.685+659C>T | 0.3901 | 0.4743 | 0.541 | 0.455 | 0.769 | 0.31084 | 0.381 |
| NM_000062.2:c.685+1100C>T | 0.2253 | 0.1124 | 0.147 | 0.262 | 0.1208 | 0.28765 | 0.274 |
| NM_000062.2:c.685+1550G>T | 0.2251 | 0.1127 | 0.152 | 0.264 | 0.1207 | 0.28734 | 0.269 |
| NM_000062.2:c.685+1770C>T | 0.2216 | 0.0992 | 0.15 | 0.262 | 0.1184 | 0.28696 | 0.27 |
| NM_000062.2:c.1029+926G>T | 0.2279 | 0.1 | 0.15 | 0.264 | 0.1224 | 0.29523 | 0.284 |
| NM_000062.2:c.1029+1443G>C | 0.2282 | 0.1004 | 0.15 | 0.269 | 0.1198 | 0.29577 | 0.284 |
| NM_000062.2:c.1029+2110T>C | 0.612 | 0.5191 | 0.469 | 0.538 | 0.2347 | 0.69271 | 0.624 |
| NM_000062.2:c.1029+2111G>A | 0.227 | 0.1003 | 0.15 | 0.264 | 0.1183 | 0.29407 | 0.283 |
| NM_000062.2:c.1030-2243T>G | 0.6129 | 0.5195 | 0.47 | 0.541 | 0.2387 | 0.69335 | 0.626 |
| NM_000062.2:c.1030-1975G>C | 0.0113 | 0.0022 | 0.008 | 0.024 | 0 | 0.01647 | 0.008 |
| NM_000062.2:c.1030-1436T>C | 0.0045 | 0.0014 | 0.001 | 0.003 | 0 | 0.00645 | 0.004 |
| NM_000062.2:c.1030-20A>G | 0.6134 | 0.5197 | 0.472 | 0.541 | 0.2461 | 0.69353 | 0.623 |
| NM_000062.2:c.1438G>A | 0.2282 | 0.1007 | 0.15 | 0.269 | 0.1202 | 0.29561 | 0.285 |
| NM_000062.2:c.*1323G>A | 0.2283 | 0.1009 | 0.151 | 0.269 | 0.1175 | 0.29578 | 0.285 |
| NM_000062.2:c.*1521G>T | 0.1496 | 0.0855 | 0.166 |  | 0.0942 |  |  |
| NM_000062.2:c.*2614A>T | 0.6058 | 0.4936 | 0.463 | 0.538 | 0.2277 | 0.69504 | 0.626 |
| PABPC4 |  |  |  |  |  |  |  |
| NM_003819.3:c.-5600T>C | 0.8127 | 0.5571 | 0.918 | 0.945 | 0.9909 | 0.90666 | 0.891 |
| NM_003819.3:c.-4432G>A | 0.0403 | <u>0.0093</u> | <u>0.095</u> | 0.024 | <u>0.3712</u> | 0.02506 | 0.047 |
| NM_003819.3:c.-4428A>G | 0.0432 | <u>0.0096</u> | <u>0.1</u> | 0.024 | <u>0.3712</u> | 0.02918 | 0.052 |
| NM_003819.3:c.-3677T>G | 0.1792 | <u>0.0535</u> | 0.122 | 0.247 | 0.1111 | 0.24258 | 0.219 |
| NM_003819.3:c.-3636G>A | 0.0052 | 0.0027 | 0.004 | 0.007 | 0 | 0.0068 | 0.007 |
| NM_003819.3:c.-3198T>C | 0.0025 | 0.001 | 0.002 | 0 | 0 | 0.00329 | 0.004 |
| NM_003819.3:c.-2286T>G | 0.0136 | 0.0031 | 0.014 | 0.01 | 0 | 0.01952 | 0.015 |
| NM_003819.3:c.-650C>T | 0.0079 | 0.0027 | 0.002 | 0 | 0 | 0.01172 | 0.008 |
| NM_003819.3:c.193+796C>G | 0.8013 | <u>0.509</u> | 0.913 | 0.945 | 0.9904 | 0.90714 | 0.895 |
| NM_003819.3:c.504-254C>A | 0.1438 | <u>0.0321</u> | 0.079 | 0.2 | 0.1093 | 0.19797 | 0.183 |
| NM_003819.3:c.738+85T>C | 0.0573 | <u>0.1955</u> | 0.012 | 0.01 | <u>0.0013</u> | <u>0.00354</u> | 0.014 |
| NM_003819.3:c.877-387C>T | 0.113 | 0.0243 | 0.065 | 0.131 | 0.1086 | 0.15452 | 0.146 |
| NM_003819.3:c.972+53A>T | 0.0018 | 0.0065 | 0 | 0 | 0 | 0 | 0 |
| NM_003819.3:c.972+704C>G | 0.0025 | 0.001 | 0.002 | 0 | 0 | 0.00328 | 0.004 |
| NM_003819.3:c.1333+26C>G | 0.0006 | 0.0001 | 0 | 0 | 0 | 0.00095 | 0 |
| NM_003819.3:c.1621-348C>G | 0.0003 | 0.0001 | 0 | 0 | 0 | 0.00042 | 0 |
| NM_003819.3:c.*765C>A | 0.0403 | <u>0.0086</u> | <u>0.096</u> | 0.024 | 0.3656 | 0.02561 | 0.049 |
| NM_003819.3:c.*1261C>T | 0.0073 | 0.0256 | 0 | 0 | 0.0006 | 0.00005 | 0.003 |
| NM_003819.3:c.*4685A>G | 0.7986 | <u>0.4999</u> | 0.911 | 0.945 | 0.991 | 0.90696 | 0.894 |
| NM_003819.3:c.*5316C>T | 0.007 | 0.0028 | 0.008 | 0 | 0 | 0.00955 | 0.008 |

*Population frequencies are taken from dbSNP. Populations with greater distance from global distribution are underlined.*

### Characterization of synonymous and missense variants of coagulation genes of interest

We found all synonymous (Supplementary Table S3) and missense (Supplementary Table S4) variants of *VKORC1, SERPING1* and *PABPC4* genes [47] from NCBI’s Single Nucleotide Polymorphism Database (dbSNP) [48] and characterized them in terms of (i) population prevalence in the Genome Aggregation Database (gnomAD) [49] [50], (ii) the percent of sequences matching the WT at that position in a multiple sequence alignment (MSA) [51], (iii) likelihood of the variant in the column of an MSA, (iv) mRNA MFE computed by both Kinefold and mFold, (v) relative synonymous codon usage (RSCU) and (vi) relative synonymous codon pair usage (RSCPU) [52] [53], (vii) rare codon enrichment [54], (viii) and %MinMax codon usage [55]. For nonsynonymous variants, we additionally used amino acid fraction matching in an MSA, likelihood of the variant amino acid in an amino acid MSA, SIFT [56] [50], and Polyphen [57] [50]. The fraction matching and MSA likelihood measures use sequence homology and may imply selection against the variant. SIFT uses sequence homology as well as physical properties of amino acids, while Polyphen uses multiple sequence and structural features to predict the effect of amino acid substitutions. MFE of mRNA may affect stability of mRNA transcripts, which will affect transcript abundance and translation. Codon and codon pair usage have been shown to impact translation kinetics [58] [59], and their metrics may be useful in assessing the impact of synonymous mutations on protein conformation and function [52]. For all variants, we provide the corresponding identifier in dbSNP (rsid) [48].

We applied filters based on codon usage changes, mRNA MFE changes, and position conservation to identify variants that were potentially impactful on protein expression or conformation, which may affect interactions with SARS-CoV-2 proteins. Then, based on population frequencies, we computed the probability of the presence of at least one filtered variant in each population, and compared with the overall probability.

For a summary of the meaning, use, and range of all scoring tools, see Supplementary Table S5.

## Results

### Computational verification of SARS-CoV-2 viral protein interactions

To study the role of coagulation in COVID-19 pathogenesis, we explored the interactions of VKORC1, SERPING1 and PABPC4 with viral proteins through computational docking. VKORC1 and ORF7a were confirmed to have strong binding affinity. Interactions are generally limited to transmembrane helices as opposed to intervening loops where warfarin is known to bind [60]. The top scoring complexes are shown in Fig 1. Plots of interface energy in Rosetta energy units against interface root mean square error for the RosettaDock results are given in Fig 2. The plots show convergence toward the minimum energy state. However, regarding SERPING1 and PABPC4, due to the lack of structural data from similar proteins, sections of the models for PABPC4 and SERPING1 were of low quality. Therefore, we continued our analysis by examining sequence homology of SARS-CoV-2 proteins to SARS proteins. Predictably, the homology was high (Table 1), suggesting that homologous SARS-CoV-2 proteins maintain interactions with human proteins as observed for SARS proteins. Specifically, several SARS proteins were found to interact with SERPING1, so it is likely that SARS-CoV-2 proteins interact with SERPING1 too. In addition, PABPC4 was found experimentally to bind to SARS-CoV-2 N protein [61].

**Fig 1.**
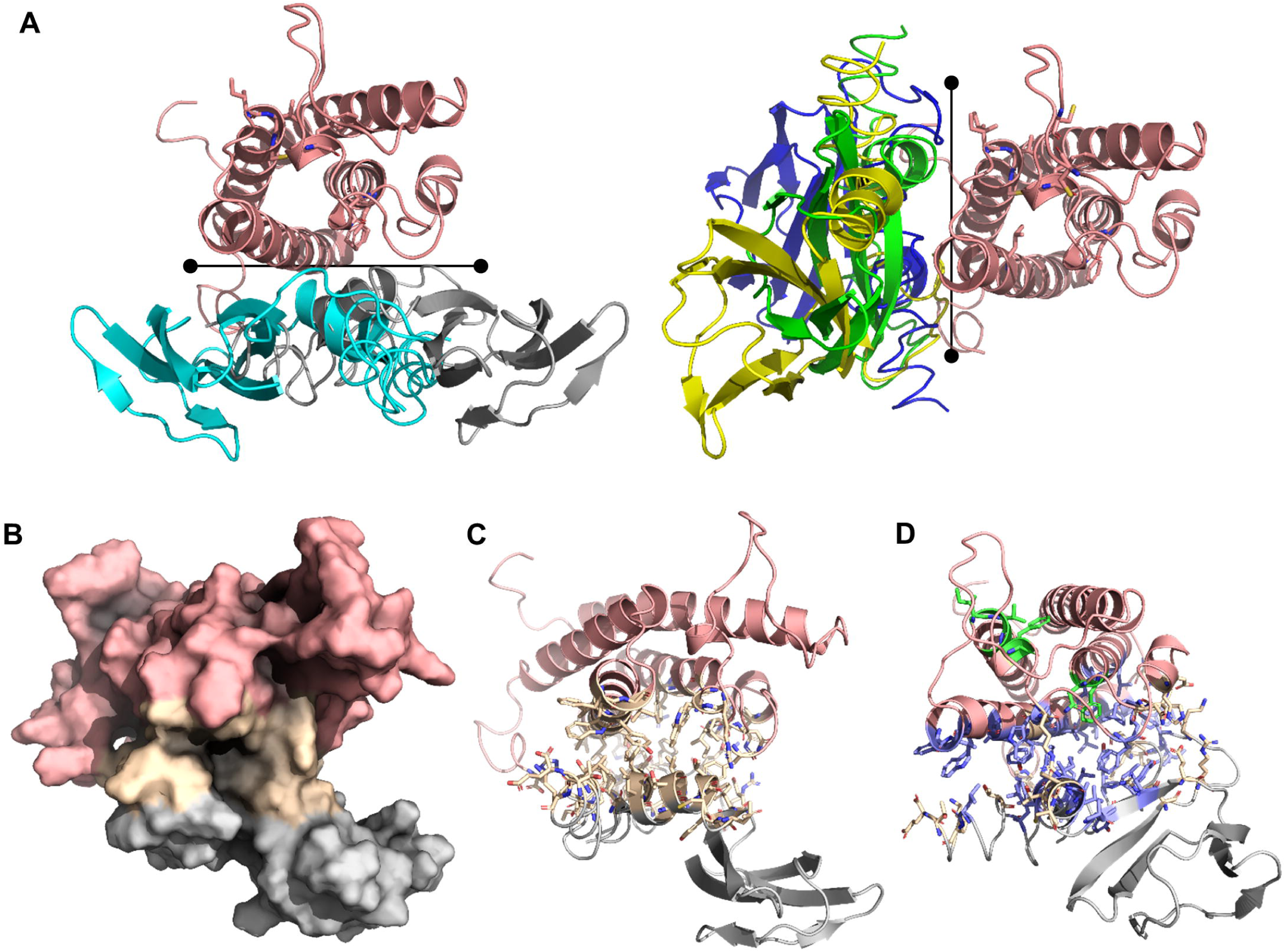
Predicted dock of VKORC1 and ORF7a proteins. A. Five protein-protein docks depict 2 binding sites (teal, grey, yellow, green, blue). B. The lowest interface-energy model is shown as a surface representation. C. The lowest interface-energy model, with side chains shown in wheat for amino acids at the interface. D. Another view of the lowest interface-energy model, with side chains shown in wheat at the interface and hydrophobics shown in blue. Amino acids of VKORC1 necessary for vitamin K binding (83F, 80N, 135C, 55F) or warfarin binding (134V, 133I) are given in green.

**Fig 2:**
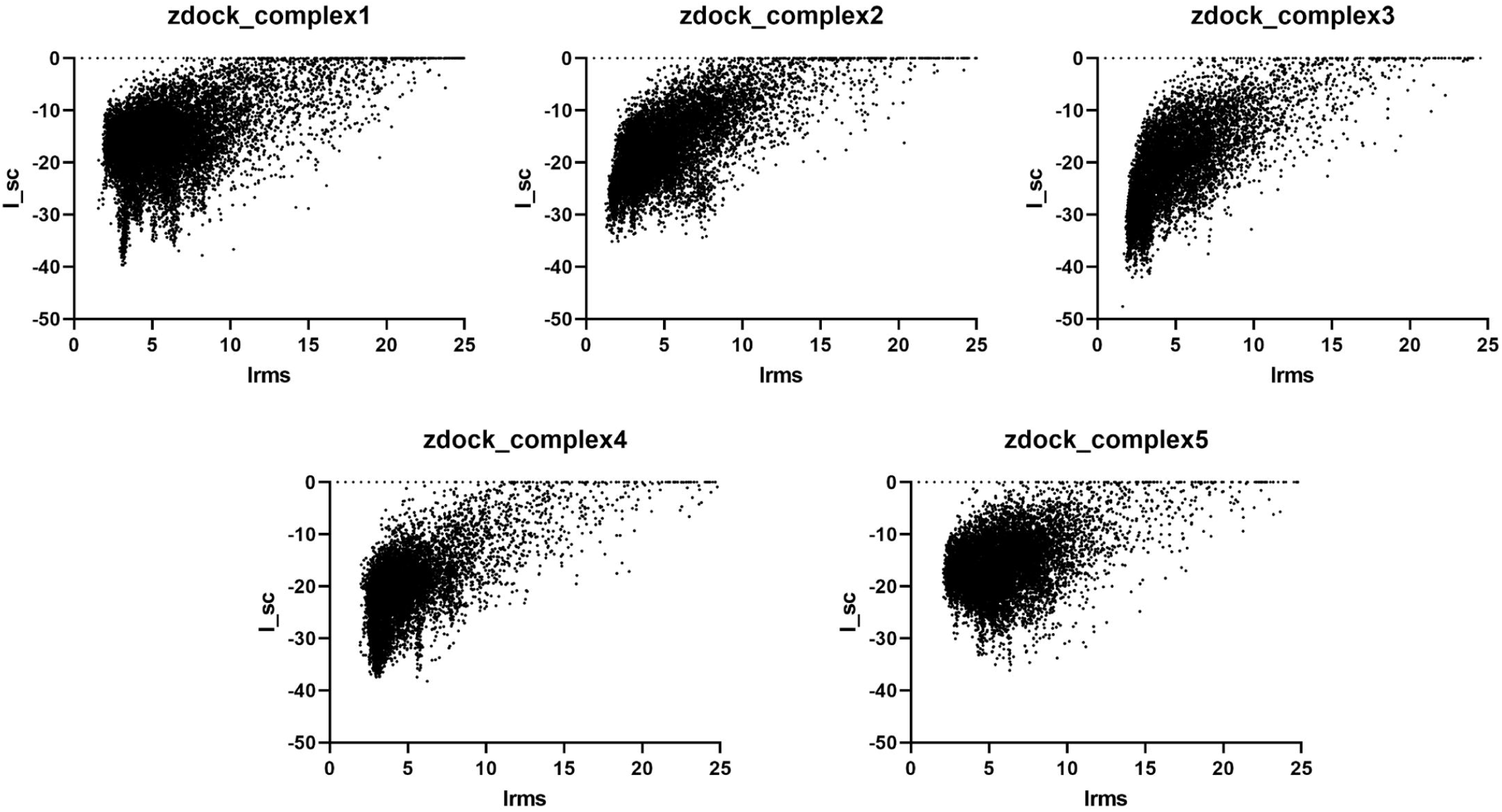
Plots of interface energy (I_sc) against interface root mean square error (I_rms). Each point represents a complex formed from one of the top 5 ZDock outputs of VKORC1 and ORF7a proteins, using 10,000 decoys. All plots form energy funnels.

### Variants that may impact COVID-19 severity

GWAS metastudies on COVID-19 outcomes recently became available [31] [32]. We focused on the impact of VKORC1, SERPING1, and PABPC4 gene variants on COVID-19 severity. While over 700 variants from these genes were found in the studies, only 55 variants had a p-value less than 0.05; these are listed in Table 2 and Table 3. However, none of them are significantly impactful when controlling for multiple hypothesis testing. Interestingly, only one of these is a coding variant. We characterized the 55 variants in terms of miRNA binding, splicing, mRNA minimum free energy, and sequence conservation, to understand how they may affect disease outcomes. miRNA’s are involved in post-transcriptional regulation by binding to mRNA transcripts, resulting in degradation of the mRNA or less efficient translation. Therefore, higher binding will most likely result in lower expressing protein. Summaries of miRNA changes are given in Table 2 and Supplementary Table S1, and full data is given in Supplementary Table S2. Interestingly, for variants which effected a change in miRNA binding potential, most caused a reduction in miRNA binding potential, which may increase protein expression. The mean change between variant and wild type miRNA affinity predictions is −11.72414, and the median is −1.

Splicing is involved in the production of mature mRNA’s for many genes. Changes in splicing may produce alternative mature mRNA’s, preventing accurate translation, and thus resulting in a protein with altered potency or affinity to the virus. While we consider splicing dysregulation as potentially impacting gene expression and disease outcome, it has rarely been shown experimentally. In vitro testing of some of these variants did not reveal differences between the splice forms and WT or substantial differences in expression. For example, Wang et al [62] examined the VKORC1 polymorphisms −1639G>A (rs9923231), 1173C>T (rs9934438), and c.-4931C>T (rs7196161) in various cell lines and did not detect any differences in expression levels. We found several intronic variants in all three genes which resulted in large changes in splicing potential (Table 2). Of these, NM_000062.2:c.52-696C>T is more common in East Asian populations, NM_000062.2:c.1029+2110T>C is more common in European populations, and NM_000062.2:c.685+1391C>T and NM_003819.3:c.1333+26C>G are comparatively rare globally.

Finally, sequence conservation gives an evolutionary view of the significance of any position in a sequence, but it is dependent on the conservation model and the quality of sequence and structural data. Several PABPC4 variants show perfect conservation at the variant position. The full data are given in Supplementary Table S1.

We found several upstream variants in VKORC1 that resulted in higher predicted miRNA binding affinity, suggesting lower expression of the protein. Of these, NM_000062.2:c.-1675G>A is relatively common in all populations (9.37% MAF). We also found several upstream variants in PABPC4 that resulted in lower predicted miRNA binding affinity suggesting higher expression of the protein.

mRNA molecules will form secondary structures based on nucleotide arrangement and affinity, which impact its structural stability. We found several variants resulting in large changes in mRNA stability. For example, NM_000062.2:c.685+1100C>T, NM_000062.2:c.1030-1975G>C, and NM_003819.3:c.*765C>A are all strongly predicted to destabilize their respective mRNA transcripts. Higher MFE may suggest higher possibility for mRNA degradation, which leads to decreased availability of transcripts and lower expression. These variants may increase mRNA degradation, reducing protein expression.

In addition, known clinical consequences of these variants are summarized in Table 4.

**Table 4:** Variations’ clinical impact.

| Variation | Clinical Impact based on Literature |
| --- | --- |
| **VKORC1** |  |
| NM_024006.4:<br>c.283+837T>C | South Indians carrying the C nucleotide require lower warfarin dosages relative to WT (T) [63]. |
| NM_024006.4:<br>c.283+124G>C | European Americans carrying the G nucleotide require lower warfarin dosages relative to WT (C) [64] |
| NM_024006.4:<br>c.174-136C>T | Turkish carrying the T nucleotide require lower warfarin dosages [65]; African Americans and European Americans carrying T nucleotide require lower warfarin dosages relative to WT (C) [66]. |
| NM_024006.4:<br>c.-1639G>A | Chinese carrying the A nucleotide require lower warfarin dosages relative to WT (G) [67]. |
| NM_024006.4:<br>c.-4931C>T | South Indians carrying the T nucleotide require increased warfarin dosages relative to WT (C) [68]. |
| **SERPING1** |  |
| NM_000062.2:<br>c.52-130C>T | Patients carrying the T nucleotide depicted worsened progression for age-related macular degeneration relative to WT (C) [69]; Chinese and Japanese carrying the T nucleotide lack an association with age-related macular degeneration, seen in Caucasian population studies, although was predicted as pathogenic [70]. |
| NM_000062.2:<br>c.1029+2110T>C | European and Mediterranean patients carrying the C nucleotide did not depict a higher association with hereditary angioedema relative to WT [71]. |
| NM_000062.2:<br>c.1030-1975G>C | The intronic polymorphism 1030 +1975G>C has no pathogenic influence on hereditary angioedema although predicted as pathogenic [71]. |
| NM_000062.2:<br>c.1030-20A>G | Association of the G allele with age-related macular degeneration was predicted to decrease the variant splicing form SERPING1, decrease protein expression and potentially limit the regulation of the compliment system [72]. No association was observed for Chinese Han carrying the G nucleotide with age-related macular degeneration [73]. |
| NM_000062.2:<br>c.-2415G>A | Chinese Han patients carrying the A nucleotide did not demonstrate an increased risk of polypoidal choroidal vasculopathy relative to WT (G) [74]. South Korean patients carrying the A nucleotide did not show association with an increased risk of leukemia relative to WT (G) [75]. Caucasians carrying the A nucleotide did not exhibit an increased risk of age-related macular degeneration relative to WT (G) [76]. |
| NM_000062.2:<br>c.52-696C>T | Patients carrying the T nucleotide did not display an increased risk for anterior uveitis relative to WT (C) [77]. Chinese Han carrying the T nucleotide did not display an increased risk for polypoidal choroidal vasculopathy relative to WT (C) [74]. Caucasians carrying the T nucleotide did not display an increased risk for age-related macular degeneration relative to WT (C) [76]. Chinese carrying the T nucleotide did not display an increased risk for diabetic retinopathy relative to WT (C) [78]. European and Mediterranean's carrying the T nucleotide did not display an increased risk for hereditary angioedema relative to WT (C) [71]. |
| NM_000062.2:<br>c.52-130C>T | Chinese carrying the T nucleotide did not display a different association with age-related macular degeneration relative to WT (C) [73]. Caucasians carrying |
|  | the T nucleotide displayed worsened progression of age-related macular degeneration relative to WT (C) [79]. Patients carrying the T nucleotide depicted worsened symptoms of age-related macular degeneration relative to WT (C) [69]. Chinese carrying the T nucleotide responded poorer to anti-VEGF treatment relative to WT (C) [80]. |
| NM_000062.2:<br>c.685+659C>T | Caucasians carrying the A nucleotide failed to depict a greater association with AMD relative to WT (G) [76]. South Korean patients carrying the A nucleotide did not depict a greater association with leukemia relative to WT (G) [81]. Han Chinese carrying the A nucleotide did not depict a significantly greater association with age-related macular degeneration relative to WT (G) [74]. |
| NM_000062.2:<br>c.685+1100C>T | European and Mediterranean patients carrying the T nucleotide failed to show a greater association with hereditary angioedema relative to WT (C) [71]. |
| NM_000062.2:<br>c.1029+926G>T | European and Mediterranean patients carrying the T nucleotide failed to show a greater association with hereditary angioedema relative to WT (G) [71]. Chinese Han carrying the T nucleotide did not depict a greater association with polypoidal choroidal vasculopathy relative to WT (G) [74]. |
| NM_000062.2:<br>c.1029+1443G>C | European and Mediterranean patients carrying the C nucleotide failed to show a greater association with hereditary angioedema relative to WT (G) [71]. |
| NM_000062.2:<br>c.1029+2111G>A | European and Mediterranean patients carrying the A nucleotide failed to show a greater association with hereditary angioedema relative to WT (G) [71]. |
| NM_000062.2:<br>c.1438G>A | Patients carrying the A nucleotide did not depict a change in Tacrolimus dosage requirements for transplant operations relative to WT (G) [82]. Chinese Han patients carrying the A nucleotide did not show a higher association with age-related macular degeneration or polypoidal choroidal vasculopathy relative to WT (G) [73]. |
| **PABPC4** |  |
| NM_003819.3:<br>c.504-254C>A | Increased risk for type 2 diabetes with the 40035928G>T polymorphism based on GWAS studies [83]. |

### Prevalence of VKORC1 variants across populations

COVID-19 has spread to the entire world, affecting people with variable genetic and racial backgrounds. Therefore, we explored ORF7a interactions with variants of VKORC1 found across races. There are 160 missense VKORC1 variants in dbSNP and at least 27 which affect warfarin sensitivity [84]. The most common variants are shown in Table 5. The locations of the warfarin sensitive variation are shown in Fig 3. However, many warfarin resistance-causing variants are not listed in dbSNP, and some do not include population frequency information. In addition, there are several intronic, upstream and downstream variants which impact warfarin dosage [85]. For example, rs9923231 (c.-1639G>A, NG_011564.1:g.3588G>A), which causes warfarin sensitivity, is very common in East Asian populations (89.95%) and comparatively less common in African populations (10.09%), with intermediate frequency for other populations.

**Table 5:**
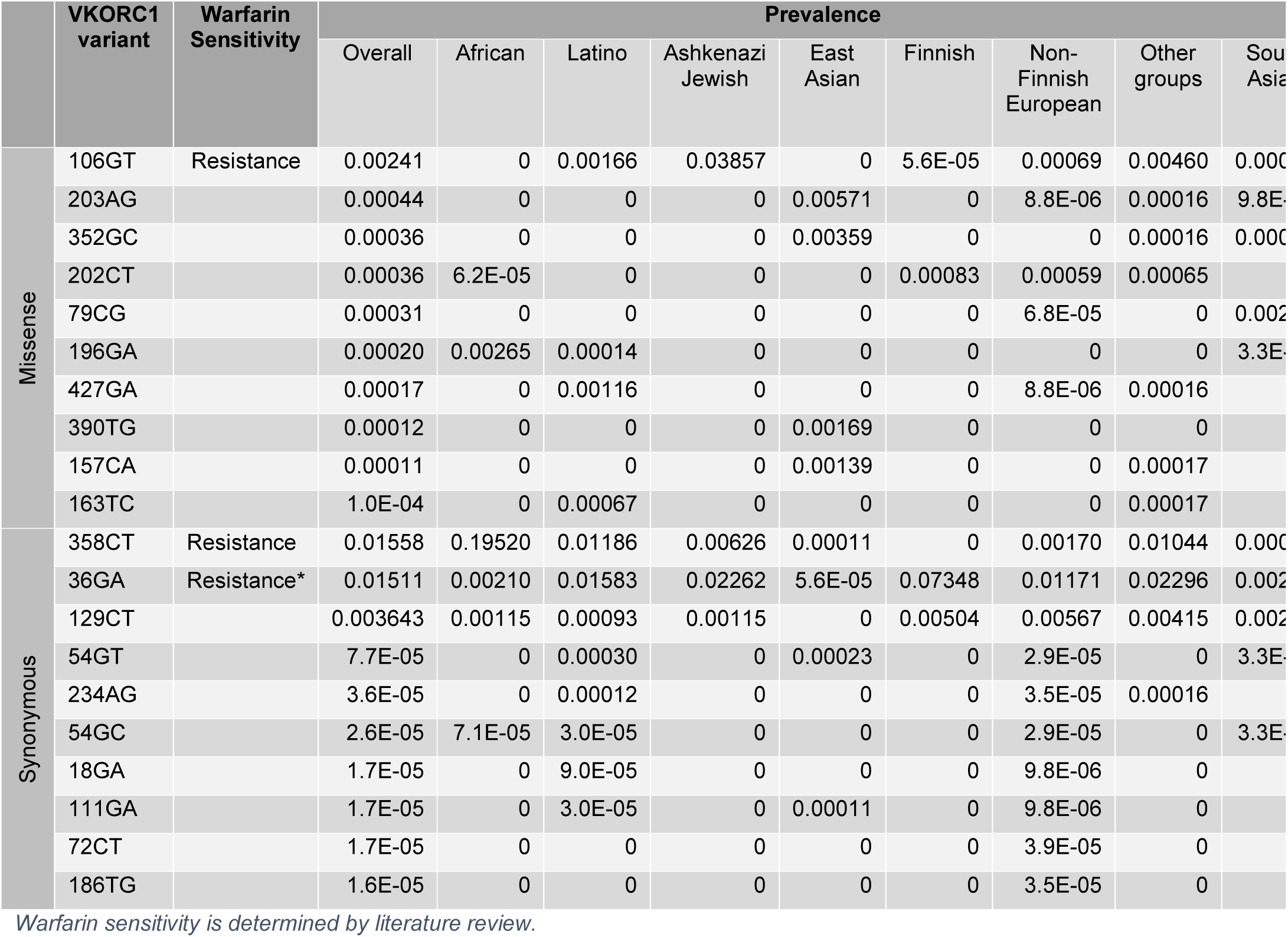
Population frequencies of missense and synonymous VKORC1 variants.

**Fig 3:**
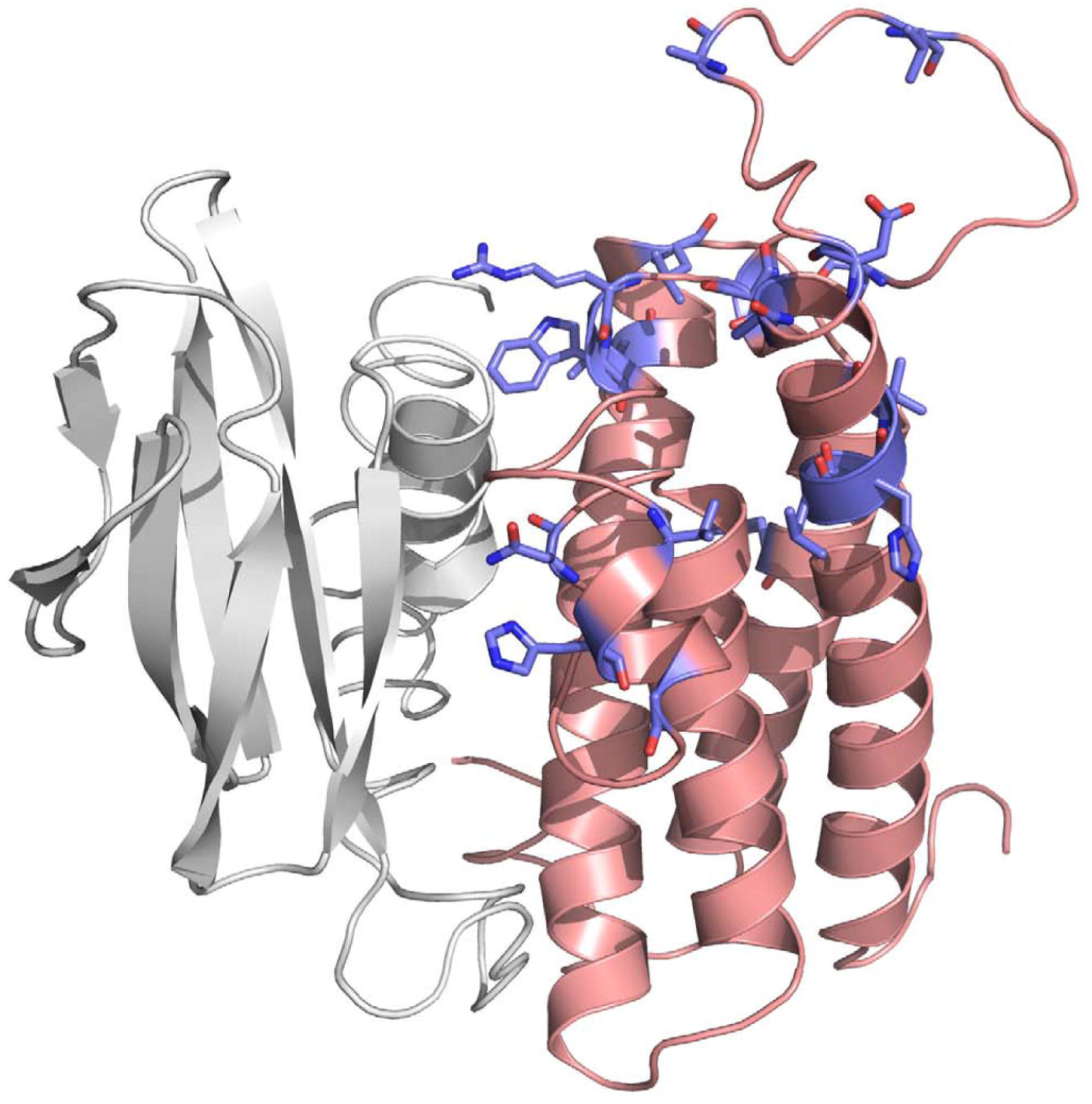
Locations of warfarin dosage affecting nonsynonymous variants in VKORC1. VKORC1 is shown in salmon, while ORF7a is shown in grey. Warfarin dosage affecting nonsynonymous variants are shown in blue.

In the United States, COVID-19 has disproportionally affected African American populations. We sought to investigate whether VKORC1 variants could be implicated in the susceptibility of this population. We found that African and African American populations were much more likely to have at least one synonymous variant that significantly changes codon and codon pair usage in a relatively conserved position. Upon further investigation, we find that this is due to a single synonymous variant, VKORC1:c.358C>T, which is very common in African and African American populations (19.52%) while comparatively rare elsewhere (maximum 1.19% among other populations). This variant is in a relatively conserved position enriched in common codons, with negative changes in relative synonymous codon and codon pair usage (RSCU, RSCPU). This variant was not predicted to change mRNA MFE, while it showed mixed results for splicing effects: while hexamer splicing scoring tools and ESEfinder showed changes in splicing near the variant, FAS ESS and Exonscan found no changes. Furthermore, this variant is associated with warfarin resistance [86] [87] [88], and in linkage disequilibrium with another variant upstream of the coding sequence (CDS), NG_011564.1:g.3350A>G, which is also common in African and African American populations (36.40%) and associated with warfarin resistance. In addition, we identified one nonsynonymous variant, VKORC1:c.106G>T, which is relatively common in Ashkenazi Jewish populations (3.857%) and rare in other populations (max 0.4599% among other populations). This variant is predicted to be deleterious by both SIFT and Polyphen and associated with warfarin resistance. This variant appears at the end of a transmembrane helix near a loop, and likely impacts loop conformation near the warfarin binding site.

These two variants were interesting, primarily due to their significant population skew. There are many other variants with different prevalence in different populations, but all others are much rarer or much more common across all populations.

We additionally characterized the population prevalence of the variants identified from the GWAS studies, finding great variance in prevalence for some. For example, NM_024006.4:c.283+837T>C is very common in all populations (64.3% MAF globally), but less common in East Asian populations (10.17%).

Furthermore, some nonsynonymous variants were identified from literature to impact drug response or disease status. Associated nucleotide changes are not always given for these variants, so characterizing them has not been possible.

### VKORC1 paralog and variants that are impactful on warfarin dosage

VKORC1L1 is a VKORC1 paralog with similar function but reduced warfarin sensitivity [89] [90]. We aligned VKORC1 with VKORC1L1 and analyzed the differences between them in the positions of variants, for additional insight into their impact on warfarin sensitivity and possible binding to ORF7a.

In the alignment of VKORC1 and VKORC1L2, seven out of twenty positions for the nonsynonymous variants impacting warfarin dosage are not conserved. This is unsurprising because the non-conserved variants are localized to the loop between transmembrane helices one and two, which is near the warfarin binding site (Fig 3). Swapping this region between VKORC1 and VKORC1L1 causes warfarin resistance in VKORC1 and warfarin sensitivity in VKORC1L1 [89].

In addition, we examined similarities of ORF7a with VKORC1 interacting proteins. Two human proteins are structurally similar to ORF7a and interact with VKORC1: CXADR, a Coxsackievirus and Adenovirus receptor [91], and PCDH1, a Hantavirus receptor [92]. Both proteins are involved in cell-cell adhesion. The structural similarity of ORF7a protein, CXADR, and PCDH1 additionally supports the interaction of ORF7a and VKORC1. The structurally aligned regions are shown in Fig 4. Structural overlap is limited to the beta sheets, with small potential for biomimicry.

**Fig 4:**
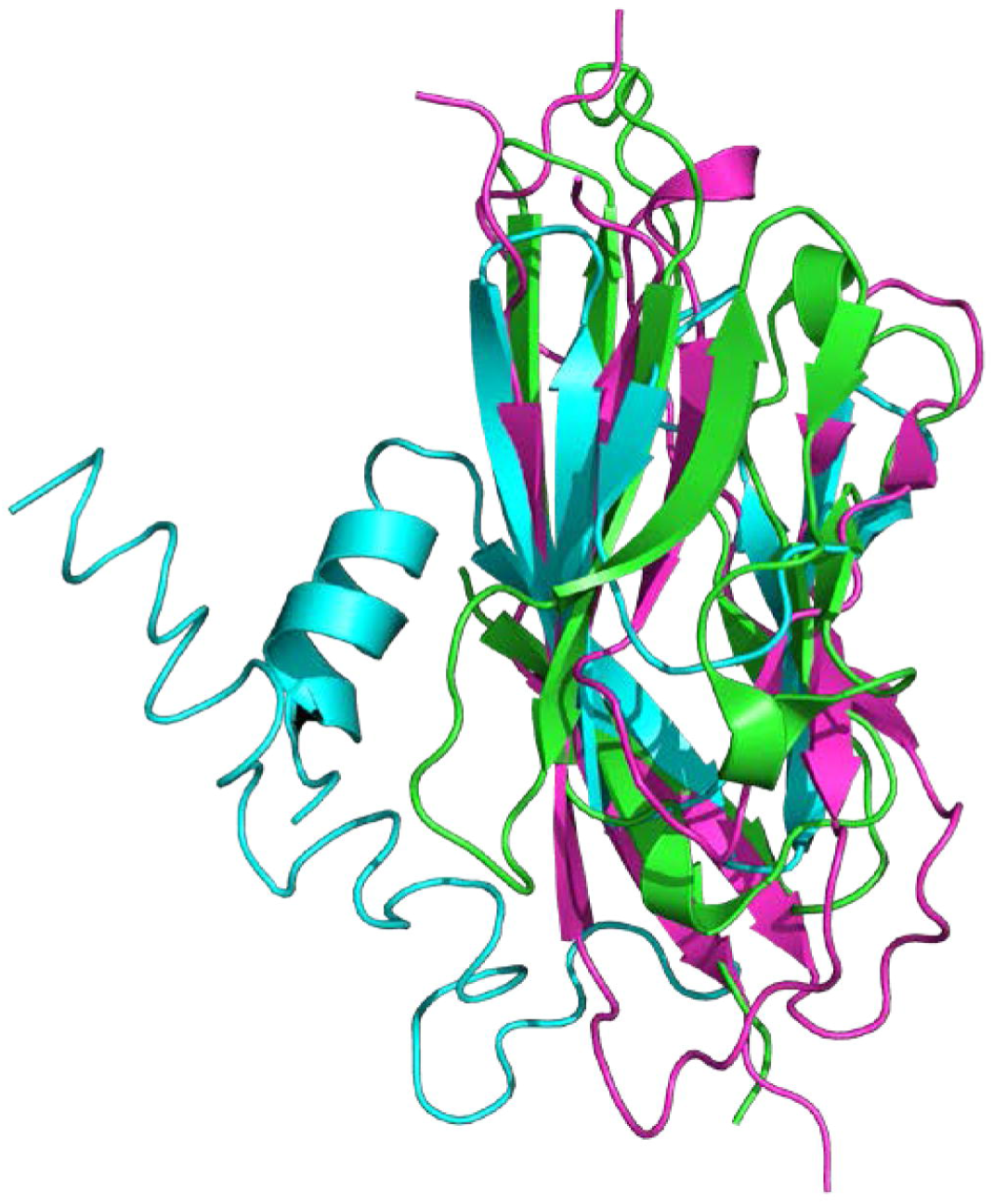
Structural alignment of ORF7a, CXADR, and PCDH1 proteins. The alignment is largely confined to the beta sheets.

## Discussion

COVID-19 is characterized by a prothrombotic phenotype of unknown etiology [1] [2] [3] [4] [6]. Developing effective treatments will require a thorough understanding of the root causes of increased coagulation that is seen in some patients. Although several possible mechanisms have been proposed [93] [94] [95] to explain why COVID-19 patients develop life-threatening clots, there may still be many parameters that have not been explored. We have identified three proteins, VKORC1, SERPING1 and PABPC4, which are related to coagulation and have been shown to interact with SARS proteins. While Pfefferle et al. (2011) focused on the viral interaction with proteins involved in the immune response, we focused instead on coagulation. We investigated computationally the binding of these proteins to SARS-CoV-2 proteins. Additionally, we identified variants of their genes and examined their prevalence across populations and their association to COVID-19. We explored mechanisms by which these variants may impact COVID-19 and we concluded that each of these proteins may provide a potential link between COVID-19 and coagulation. Furthermore, we identified the gaps in knowledge that need to be addressed to further explore their roles.

VKORC1 is crucial for maintaining active vitamin K levels and hence for the function of several essential coagulation factors. We generated strong computational evidence pointing to an interaction between VKORC1 and ORF7a that are in agreement with previous experimental data. However, the impact of this interaction is currently unknown. Given the prothrombotic phenotype that is seen in severe COVID-19 patients, we hypothesize that binding of ORF7a to VKORC1 has an effect on coagulation that may be altered in patients with certain VKORC1 polymorphisms.

Reduced vitamin K levels were suggested to be related to worse prognosis in COVID-19 patients [96]. This may be due to the role of vitamin K in the production of coagulation factors and coagulation regulatory proteins, the inverse relationship of vitamin K and inflammatory response [97] [98], or the inverse relationship of vitamin K and Interleukin-6 (IL-6) levels [99]. The inflammatory and immune response is a significant cause of symptoms in COVID-19 patients [100]. These factors are likely linked: inflammation causes production of IL-6, which initiates coagulation activity [101], exhausting the supply of coagulation factors and active vitamin K. Patients exhibiting warfarin resistance, for example patients with the 106G>T missense variant, may produce more active vitamin K, increasing the available supply of coagulation factors and producing more severe clotting, possibly even DIC. Alternatively, the presence of variants may lead to altered conformation leading to differential binding to either warfarin or ORF7a. Interestingly, synonymous variant 358C>T is characterized by a large change both in RSCPU and RSCU suggesting that it may be associated with altered cotranslational folding. The inflammation resulting from COVID-19 infection is of particular interest, as children are experiencing increased rates of Kawasaki disease-like symptoms [102], especially in areas where COVID-19 outbreaks were worse.

In addition, the VKROC1 - ORF7a interaction may also have an impact on tetherin function. SARS ORF7a is known to inhibit tetherin [103] [104], also known as BST-2. Tetherin inhibits virion dispersal [105], and several viruses, including HIV, have auxiliary proteins to counter this effect. The structures of tetherin and VKORC1 are noticeably similar, sharing a coiled-coil architecture: VKORC1 has four alpha helices bound together [60], while tetherin exists as a homomer of four alpha helices [106] [107]. Reduced VKORC1 expression or binding to ORF7a, as it may occur with some variants, may increase the availability of ORF7a to bind and inhibit tetherin increasing the severity of SARS-CoV-2 infection. Of note, ORF7a has the highest RSCU of any SARS-CoV-2 protein [108]. This may make it efficiently translated, resulting in high expression levels to more effectively counter the effect of tetherin.

Due to the limited availability of homologous structural data, homology models of PABPC4 and SERPING1 could not be constructed with high confidence. Therefore, it wasn’t possible to create complexes to model and analyze the interactions between these proteins and viral proteins. Experimental validation would be instrumental in confirming these interactions. While PABPC4 has been found to interact with SARS-CoV-2 N protein experimentally [61], interactions between VKORC1 and ORF7a have not been confirmed. Similarly, interactions between SERPING1 and viral proteins have not yet been tested but would help inform our hypothesis.

COVID-19 has had an unequal impact on populations across the globe [109] [110]. In the United States, as elsewhere, some populations are proving to be more susceptible to the disease and its complications. A large number of factors influence the presentation and prognosis within populations, including age, access to health care, and presence of comorbidities such as diabetes, hypertension, advanced age, and chronic lung disease [111] [112]. Understanding the relative risks and associated complications of COVID-19 on a population level provides valuable information to help clinicians plan the most beneficial course of treatment. Unfortunately, detailed demographic data, including breakdown by age, race, gender, and comorbidities, is incomplete; as of May 30, 2020, only 22% of reported cases have information on reported comorbidities and only 45% of reported cases nationwide were presented with demographic data sufficient to determine race [112]. In addition, there is increasing evidence that African-American populations are at higher risk of thrombotic events with COVID-19, even when adjusting for common risk factors such as BMI, diabetes, hypertension, and cardiovascular disease [113] [114]. This suggests a role for additional molecular factors that can contribute to a predisposition to thrombosis in the presence of COVID-19.

While many factors can influence the presentation of disease, the impact of host genetic variants on viral protein interaction is not clear. In host proteins interacting with viral proteins, genetic variants may affect this interaction. Reducing the strength of this interaction or the availability of host proteins may reduce the effectiveness of viral proteins and the severity of infection. On the other hand, host genetic variants may affect protein function or availability. Viral interactions may exacerbate these affects.

For example, SERPING1 is an inhibitor of plasma kallikrein, which produces bradykinin from high-molecular-weight kininogens. Genetic variants that reduce the translation efficiency of SERPING1 may reduce SERPING1 activity, and viral interactions with SERPING1 may further reduce activity, resulting in excessive levels of bradykinin and possibly angioedema.

Further complicating this situation, ACE2 inactivates des-Arg^9^ bradykinin (DABK) [115] [116], an active bradykinin metabolite, and reduced ACE2 activity is associated with enhanced signaling of DABK, angioedema, and neutrophil infiltration in the lungs [117] [115]. Reductions in ACE2 activity or availability caused by genetic variants or other conditions may be exacerbated by SARS-CoV-2 infection. The combined effect of ACE2 and SERPING1 inhibition by viral proteins may cause excessively high levels of bradykinin and fluid in the lungs [116].

There is strong evidence, both computational and experimental, of the binding of ORF7a and VKORC1. We suggest that some VKORC1 variants may affect pulmonary intravascular coagulopathy observed in COVID-19. The extensive damage and clotting in the lungs may exhaust coagulation factors and active vitamin K necessary for carboxylation of coagulation factors. The binding of ORF7a and VKORC1 may also limit coagulation in the lungs by preventing the reduction of vitamin K epoxide to active vitamin K. This interaction may be less influential in patients with warfarin resistance due to increased production of VKORC1 protein or modified VKORC1 conformation, resulting in increased coagulation and worse prognosis.

In addition, the interaction of SERPING1 and PABPC4 with viral proteins may result in dysregulated coagulation and immune response. Specifically, the combined effect of ACE2 and SERPING1 interacting with viral proteins may increase levels of bradykinin, potentially causing fluid leaking into the lungs. As with VKORC1, genetic variants may impact the efficacy of viral inhibition by changing protein conformation or expression. Because these genetic variants appear at different frequencies in different populations, this may impact outcomes for COVID-19 patients from different ethnic groups.

## Supporting information

Computed features for all genetic variants of interest in VKORC1, SERPING1, and PABPC4 from GWAS studies.

Predicted binding score for all relevant miRNA species and the variant (MUT) and wild type (WT) sequences.

Computed features for all identified synonymous variants identified in proteins that interact with SARS proteins.

Computed features for all identified missense variants identified in proteins that interact with SARS proteins.

Description, explanation and range of computed features.

## Supporting information

**S1 Table: Computed features for all genetic variants of interest in VKORC1, SERPING1, and PABPC4 from GWAS studies**.

**S2 Table: Predicted binding score for all relevant miRNA species and the variant (MUT) and wild type (WT) sequences**.

**S3 Table: Computed features for all identified synonymous variants identified in proteins that interact with SARS proteins**.

**S4 Table: Computed features for all identified missense variants identified in proteins that interact with SARS proteins**.

**S5 Table: Description, explanation and range of computed features**.

## Acknowledgements

This research was in part supported by the Intramural Research Program of the National Library of Medicine at the National Institutes of Health.

This study was in part supported by the National Institutes of Health grant HL151392.

